# Highly expressed genes evolve under strong epistasis from a proteome-wide scan in *E. coli*

**DOI:** 10.1101/185967

**Authors:** Eric Girard, Pouria Dasmeh, Adrian W.R. Serohijos

**Author notes:** Equal contribution (Eric Girard, Pouria Dasmeh).

## Abstract

Epistasis or the non-additivity of mutational effects is a major force in protein evolution, but it has not been systematically quantified at the level of a proteome. Here, we estimated the extent of epistasis for 2,382 genes in *E. coli* using several hundreds of orthologs for each gene within the class *Gammaproteobacteria*. We found that the average epistasis is ~41% across genes in the proteome and that epistasis is stronger among highly expressed genes. This trend is quantitatively explained by the prevailing model of sequence evolution based on minimizing the fitness cost of protein unfolding and aggregation. Our results highlight the coupling between selection and epistasis in the long-term evolution of a proteome.

## INTRODUCTION

Resolving the link between genotype and phenotype or the fitness landscape is a central goal in molecular biology and evolution. Knowledge of the structure of the fitness landscape will lead to a better understanding of the evolutionary origin of natural proteins and to solutions to practical evolutionary problems, from rational design of enzymes to the development of new antibiotics^1^. The fitness landscape is complex and a consequence of this complexity is *epistasis* or the dependence of mutational effects on genetic background^2^. The presence of epistasis implies that the effects of multiple mutations are non-additive and that their order of fixation matters. Indeed, epistasis directly affects the potential pathways to explore the fitness landscape^2^. Despite the many experimental and theoretical studies on detecting and elucidating its role in molecular evolution^3^,^4^, none has investigated the strength of epistasis at a proteome-wide level. Such an analysis can determine correlations between epistasis and genomics properties that could hint of a universal mechanism, if any, for epistasis in proteome evolution. Additionally, a mechanistic understanding of epistasis has practical applications; as yet, it is rarely accounted for in the molecular evolution toolboxes for quantifying from genomic sequences the strength of multiple evolutionary forces—mutation, drift and selection^4^,^5^.

To estimate the epistasis experienced by genes in long-term evolution, one approach is to compare two rates of amino acid substitutions^6^. These two rates are the average pairwise substitution rate *R*_dN/dS_, which is background-dependent, and the rate of mutational usage *R*_*u*_, which is background independent. Both rates are calculated from a multiple sequence alignment (MSA) of a gene’s orthologs. Specifically, the extent of epistasis is quantified as

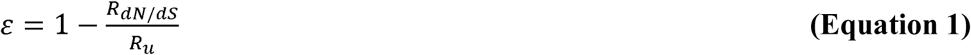

*R*_dN/dS_ is the average d*N*/d*S* (the ratio of non-synonymous substitution rate *dN* and synonymous substitution rate *dS*) for all pairs of orthologues in an MSA. *R*_dN/dS_ is calculated over the entire length of the gene. thus it reflects the co-evolution between sites. This is also implies that *R*_dN/dS_ accounts for the background-and lineage-specificity of amino acid substitutions. *R*_*u*_ = 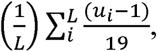 where *u* is the mutational usage and is the number of unique amino acids in each site in an MSA. *L* is the length of the protein. *R*_*u*_ represents the ratio between observed accessible amino acid substitutions in a site. (*u*-1). and all possible amino acid substitution assuming no selection. that is. (20-1) = 19. Unlike *R*_dN/dS_, *R*_u_ simply counts the number of unique amino acids per site. thus it does not reflect the co-evolution between sites in the protein. Therefore. *R*_*u*_ is independent of background and lineage. When all mutations are neutral. both *R*_*u*_ and *R*_*dN/dS*_ are equal to 1. When *random* mutations are not neutral. such as in proteins where they are predominantly destabilizing and deleterious, purifying selection will lead to *R*_*u*_ and *R*_*dN/dS*_ less than 1. However. the presence of epistasis implies that genetic background further screens substitutions. thus the the background dependent *R*_*dN/dS*_ is slower than the background independent *R*_*u*_. The expression for epistasis (**Eq. 1**) estimates how much epistasis or the background specificity of mutational effects slows down the rate of protein evolution. Kondrashov and coworkers^6^ applied this method to estimate epistasis in the long-term evolution of 16 mammalian proteins and found epistasis to vary from ~40% to ~80%. This slowing down of evolutionary rates can also arise from the heterogeneity of fitness effects of mutations^8^.

## RESULTS

### Proteome-wide espistasis in *E. coli*

To generalize this observation. we estimated epistasis in the evolution of 2.382 genes in *E. coli* using thousands of orthologs within the *gammaproteobacteria* (see Methods and Supporting Information). We calculated *R*_u_ and *R*_dN/dS_ from the multiple sequence alignment of each gene (Methods). The rates *R*_u_ and *R*_d*N*/d*S*_ and the epistasis for each gene are shown in **Fig. 1a** and their distributions in **Fig. 1b**. Since epistasis is expected to slow down the rate of evolution. the lineage-independent rate *R*_*u*_ is greater than the lineage-and background-dependent *R*_d*N*/d*S*_ (note the deviation from R_u_ = R_dN/dS_ line in **Fig. 1a**; Wilcoxon signed-rank test. p-value<10^−16^). The average *R*_u_ and *R*_d*N*/d*S*_ are 0.36 ± 0.09 and 0.20 ± 0.08. respectively. which lead to a proteome-wide epistasis of ~41% (**Fig. 1b**; full data is listed in **Table S1**). This estimate implies that epistasis and background specificity of mutational effects in proteins slows down the evolutionary rates of proteins in *E. coli*. on average. by ~41%. The magnitude of epistasis is broadly distributed with some genes experiencing epistasis of up to ~80% (**Fig. 1b**). These estimates over several thousand genes is slightly lower than the value calculated by Breen *et al.*^6^ for 16 mammalian proteins.

**Figure 1.**
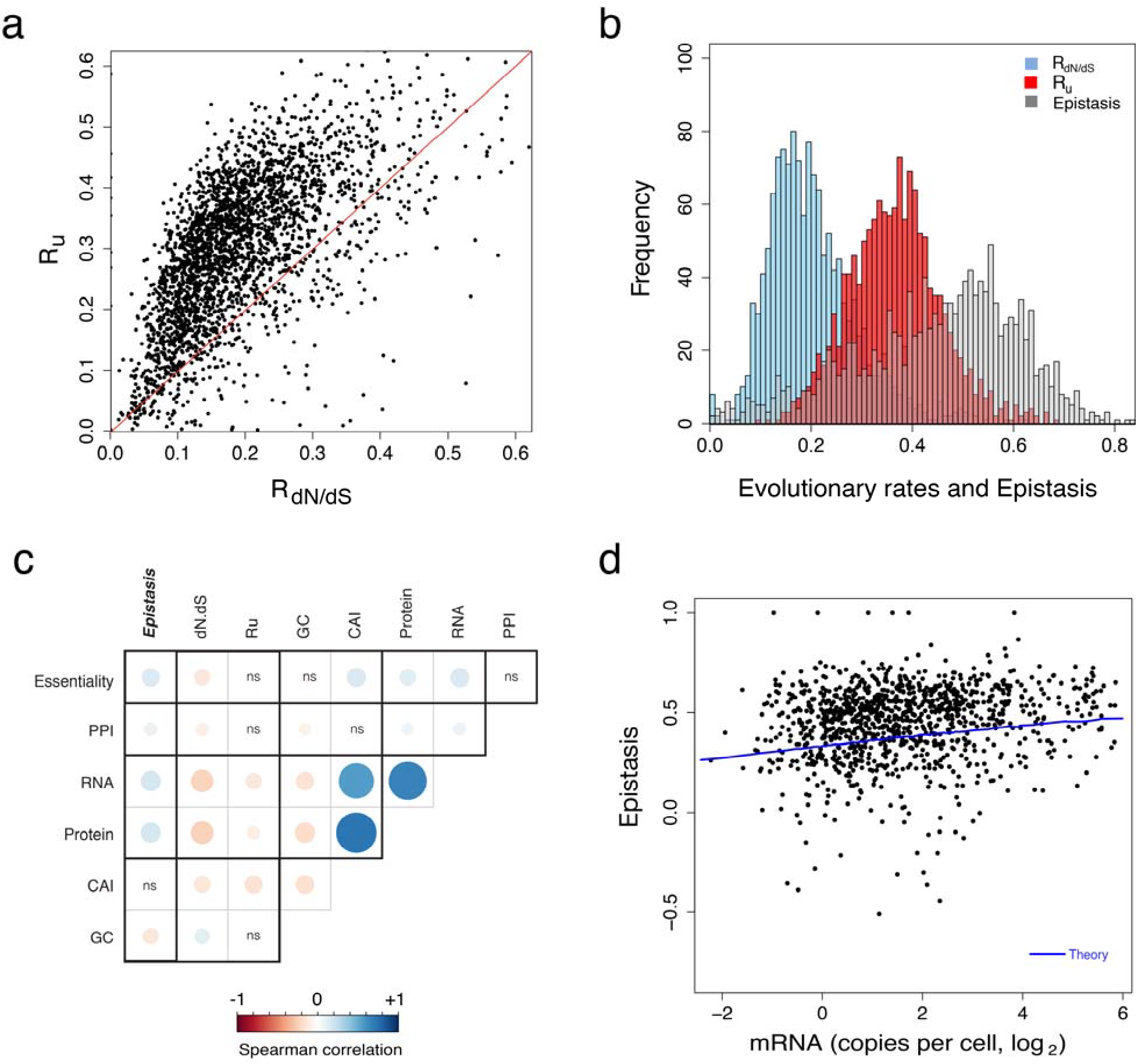
Proteome-wide estimate of epistasis in *E. coli*. **(a)** Background-dependent evolutionary rate *R*_*dN/dS*_ is significantly slower than the background-independent rate of mutational usage *R*_*u*_ (Wilcoxon signed-rank test. p-value<10^−16^). **(b)** The average epistasis is ~41 ± 16% among 2.382 genes in *E. coli*. **(c)** Epistasis is positively correlated with genome-wide factors: mRNA and protein expression levels. essentiality of proteins. number of protein-protein interactions and codon adaptation index (CAI) (see Table S4 for the correlation coefficients and p-values). Boxes labeled “ns” are not significant (p-value>0.05). **(d)** Highly expressed genes experience strong epistasis (Spearman r = +0.17. p-value<10^−9^). which can be explained by a model of sequence evolution based on selection against protein misfolding and aggregation (blue line; see also **Figs. S3, S4** & **S10**).

Next, we checked for the robustness of our results to factors that may influence the calculation of substitution rates and epistasis. First. the evolutionary rates. in particular *R*_*u*_ that count the number of unique amino acids. are sensitive to the number of orthologs in an MSA. Too few orthologs may lead to undersampling of *R*_*u*_ and to negative values for epistasis. However. as shown by the plot of epistasis versus the number of orthologs (**Fig. S1**). this artifact is present only in genes with MSA alignments less than 200 orthologs. Second. non-fixed amino acid states or polymorphisms can inflate the mutational usage *R*_*u*_ and epistasis. To estimate the impact of non-fixed states in our amino-acid usage calculation. we used a correction based on the probability of occurrence of non-fixed amino acids at given site in the alignment (**Fig. S2** and Supporting Information). The average correction to *R*_*u*_ based due to non-fixed polymorphism is only ~<u>±</u>2 % (**Fig. S2**). **Table S2** presents amino acid usage correction along with probability of observing a non-fixed state as fixed for all genes. Lastly. the calculation of *R*_d*N*/d*S*_ is sensitive to saturation and the counting method. We control for this effect by using seven counting methods (five heuristic and two maximum-likelihood codon-based approaches) for *dN*, *dS*, and *dN/dS* (**Figs. S5, S6, S7** and **Table S3**); we find no significant difference among values calculated with different counting methods. Altogether. our estimates of epistasis are robust to the number of orthologs. presence of polymorphisms. and approaches for counting substitution rates.

### Relationship of epistasis with genomic properties

The rate of protein evolution is influenced by several factors ranging from molecular. to cellular. and to population level^9^. We determined if epistasis is also influenced by these factors. Specifically. we calculated the correlation between epistasis and publicly available data on mRNA expression level. protein abundance. gene essentiality. protein-protein interaction (PPI). and codon adaptation index (CAI) (**Fig. 1c, Table S4** see Methods). We found that epistasis shows a weak yet significant positive correlation with number of orthologs. PPI. mRNA and protein expression levels. as well as CAI. As shown previously^10^. *R*_d*N*/dS_ negatively correlates with expression level (*r* = −0.24. p-value<10^−16^) implying that highly expressed genes are under strong purifying selection. Since *R*_*u*_ also reflects selection. it similarly shows negative correlation with expression level (*r* = −0.18. p-value<10^−10^). However. the weaker anti-correlation between *R*_*u*_ and expression level compared to that of *R*_d*N*/dS_ leads to a positive correlation between epistasis and expression level (*r* = +0.17. p-value<10^−9^) (**Fig. 1d**). This finding implies that background specificity significantly slows down the rate of evolution among highly expressed genes. Thus. our proteome-wide estimates demonstrate that highly expressed genes not only experience stronger purifying selection. but also greater epistasis in their long-term evolution. This result highlights the coupling of selection and epistasis in proteome evolution^11^.

### Model of sequence evolution based on protein folding explains proteome-wide correlation between epistasis and expression level

The negative correlation between evolutionary rate *R*_dN/dS_ and expression level is well-established^10^,^12^,^13^. This observation have been explained by a model of sequence evolution based on selection against protein misfolding due to mistranslation^10^ and/or genetic mutations^12^,^14^. The biological rationale is that misfolded proteins can form aggregates are toxic to the cell^15^,^16^. To determine if this same hypothesis can quantitatively explain the trend between epistasis and mRNA expression level. we combine the population genetic formalism for evolutionary rate with protein folding thermodynamics^12^,^17^^-^^19^. Assuming that cellular fitness *F* is *inversely* proportional to the total number of misfolded proteins in the cell. it may be formally written as^10^:

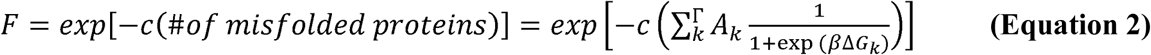

**Eq. 2** expresses the probability that the protein product of gene *k* is unfolded as a function of its stability Δ*G*_*k*_. The energy factor *β* = 1/*k*_*b*_*T* where *k*_*b*_*T* ~ 0.59 kcal/mol at room temperature. This probability multiplied by the cellular abundance of the gene *A*_*k*_ gives the number of misfolded copies (**Fig. S10**). The summation extends over all genes *Γ* in the proteome. The parameter *c* is the fitness cost of each misfolded protein (~10^−7^) (ref.^16^). As shown previously^10^,^12^ and in our specific dataset (**Figs. S3** and **S4**). this fitness function recapitulates the trend between d*N*/d*S* and expression level. But to arrive at epistasis. we also need a theoretical estimate for the mutational usage *R*_*u*_. In a recent work^17^. we showed that *R*_*u*_ is the rate of evolution of the most stable sequence in an MSA. By simulating sequence evolution (Supporting Information). we can arrive at a theoretical MSA evolved under the fitness function (**Eq. 2**) and then calculate *R*_*u*_ (**Fig. S4**).

Highly expressed (and more abundant) genes are under strong purifying selection; thus. *R*_*u*_ and *R*_*dN/dS*_ negatively correlate with mRNA level. both in theory (**Fig. S4**) and in *E. coli* (**Fig. S3**). More interestingly. the theoretical dependence of *R*_*u*_ vs. mRNA is weaker than *R*_*dN/dS*_ vs. mRNA leading to stronger epistasis among highly expressed genes (**Fig. S4**). Thus. selection against protein misfolding can explain the genomic observation that highly expressed and more abundant genes experience stronger epistasis (Fig. **1c,d**). A geometric interpretation of epistasis is the curvature of the fitness landscape; indeed. for the genotype-phenotype relationship based on folding stability (**Eq. 2**). the landscape exhibits greater curvature at higher expression levels (**Fig. S10**).

## DISCUSSION

Our study. for the first time. provides proteome-wide estimate of epistasis in *E. coli*. On average protein in E. coli evolves with ~41% epistasis. One interpretation of this result is that the rate of protein evolution is reduced by 41% due to background dependence of mutational effects. Moreover. we found that highly expressed proteins evolve with stronger epistasis. which can be explained by selection against protein misfolding. Our results highlight the coupling between selection and epistasis. which has been demonstrated in specific proteins^4^,^20^. but not in the longterm evolution of a proteome.

This work focused on the E. coli proteome; however. it will be interesting to generalize these observations in other well-studied model organisms such as yeast. worm. fly. mouse. and human. where selection (detected by d*N*/d*S*) has been shown to strongly correlate with expression level^10^. Demonstrating these results across all kingdoms of life could generalize the finding that selection due to folding stability is a universal mechanism for some of the epistasis experienced in the long-term evolution of a proteome.

## Methods

### Sequences and alignment

List of genes for *Escherichia coli* K-12 MG1655 was taken from NCBI. From this list (4140 genes), KEGG ids (total of 3059 ids) were used to retrieve functional orthologs within *Gammaproteobacteria* class. Ortholog sequences were available for 2814 of the 3059 genes. To optimize alignment, sequences 15% longer or shorter from the reference E. *coli* gene were removed from the set. DNA sequences were converted to protein sequences prior to alignment and calculation of amino-acid usage. For the protein alignments, we used default parameters except for the allowed positions with gaps that were set to half, to allow gaps at positions where less than 50% of sequences had gaps.

### Evolutionary rate analysis

The amino-acid usage measure can be used to obtain an estimation of d*N*/d*S* ratio under the assumption of non-epistatic evolution. The amino-acid usage <*u*> is defined as the number of different amino-acids observed at one site, averaged over all sites in an alignment. We can then estimate non-epistatic d*N*/d*S* from <*u*> using (*u*-1)/19 where (*u*-1) is the number of amino-acid states into which the current amino acid can be substituted, divided by 19 amino-acid possibilities. The choice of a proper and unbiased method to estimate d*N*/d*S* is crucial in the current work. We thus systematically checked performance of five different heuristic counting approaches and two maximum-likelihood (ML) codon models for 3124 genes in E.coli and concluded that the simplest model of Nei and Gojobori gives the most unbiased d*S* and d*N*/d*S* estimates which reasonably fit values from accurate yet computationally expensive ML methods. For the complete analysis check Supplementary methods and Table S4.

### Theoretical model

To calculate the extent of epistasis, we used an expression for substitution rates that takes the stability effects of mutations into account (Equation 1). Fitness is proportional to the number of misfolded copies in the cell which in turn is a product of total abundance and the probability of being in the folded state. This decomposition enables us to consider the known distribution of mutational effects on protein stability as distribution of fitness effects and calculate evolutionary rates accordingly (see Supplementary information for full analysis). We then took mRNA abundance for each protein in this study, converted mRNA abundance to protein abundance and evaluated the rate of mutational usage, *R*_*u*_ and pairwise rate of evolution, *R*_dN/dS_ as explained in Supplementary information.

## Acknowledgments

We thank members of the Serohijos lab for Discussion and Christine Angus-Banaszek for a careful reading of the manuscript. We acknowledge funds from the University of Montreal, NSERC, and the Merck Foundation.

## Competing interests

The authors declare no competing financial interests.

## Author Contribution

AS and PD conceived and designed the experiments. EG and PD performed the analyses. All authors contributed to the writing.

